# Light-dependent Control of Bacterial Expression at the mRNA Level

**DOI:** 10.1101/2022.07.30.502174

**Authors:** Américo T. Ranzani, Markus Wehrmann, Jennifer Kaiser, Marc Juraschitz, Anna M. Weber, Georg Pietruschka, Günter Mayer, Andreas Möglich

## Abstract

Sensory photoreceptors mediate numerous light-dependent adaptations across organisms. In optogenetics, photoreceptors achieve the reversible, non-invasive, and spatiotemporally precise control by light of gene expression and other cellular processes. The light-oxygen-voltage receptor PAL binds to small RNA aptamers with sequence specificity upon blue-light illumination. By embedding the responsive aptamer in the ribosome-binding sequence of genes of interest, their expression can be downregulated by light. We developed the pCrepusculo and pAurora optogenetic systems that are based on PAL and allow to down- and upregulate, respectively, bacterial gene expression using blue light. Both systems are realized as compact, single plasmids that exhibit stringent blue-light responses with low basal activity and up to several ten-fold dynamic range. As PAL exerts light-dependent control at the RNA level, it can be combined with other optogenetic circuits that generally control transcription initiation. By integrating regulatory mechanisms operating at the DNA and mRNA levels, optogenetic circuits with emergent properties can thus be devised. As a case in point, the pEnumbra setup permits to upregulate gene expression under moderate blue light whereas strong blue light shuts off expression again. Beyond providing novel signal-responsive expression systems for diverse applications in biotechnology and synthetic biology, our work also illustrates how the light-dependent PAL-aptamer interaction can be harnessed for the control and interrogation of RNA-based processes.

## Introduction

Sensory photoreceptor proteins enable organisms to sense light stimuli and translate them into physiological adaptations^1^. As one photoreceptor class, light-oxygen-voltage (LOV) receptors respond to blue light and commonly comprise modular sensor and effector domains^2^. Light absorption by a flavin-nucleotide chromophore within the LOV sensor initiates a well-characterized photocycle during which a conserved cysteine residue covalently bonds to the atom C4a of the flavin isoalloxazine heterocycle^3^. This reaction is accompanied by protonation of the neighboring flavin N5 atom which triggers hydrogen-bonding and conformational rearrangements and thereby converts the photosensory input into biochemical signal outputs^4–6^. The underlying structural transitions propagate from the LOV sensor to the effector module via linker segments that are frequently α-helical in structure. Individual LOV receptors regulate as a function of blue light different effector outputs, including kinases, transcription factors, and phosphodiesterases^1,7,8^. Returned to darkness, the covalent thioadduct ruptures in a base-catalyzed reaction^9^, denoted dark recovery, and the LOV receptor reverts to its resting state. Judiciously chosen residue exchanges close-by the chromophore greatly modulate the kinetics of dark recovery^10^.

Beyond their pivotal roles in nature, LOV and other photoreceptors have been harnessed for optogenetics^11^ to control by light cellular state and physiology upon heterologous expression in target organisms. Originally explored in the neurosciences and restricted to microbial rhodopsins^12,13^, the optogenetic principle has since been extended to other research areas and photoreceptor classes. An ever-growing set of engineered receptors with customized, light-gated activity now complement natural photoreceptors^14,15^. Together, these receptors allow the spatiotemporally precise and reversible control of diverse biological processes, including protein-protein and protein-DNA interactions, transcription, recombination, epigenetics, subcellular localization, cytoskeleton dynamics, and signaling cascades. Despite these advances, sensory photoreceptors that directly bind to and act on RNA in light-dependent manner had long been lacking. In addition to the long-known mRNAs, rRNAs, and tRNAs that are central to transcription and translation, multiple, more recently discovered RNA species govern diverse aspects of gene expression and other cellular processes. Against this backdrop, the description of the LOV receptor PAL from *Nakamurella multipartita* as a light-regulated RNA-binding protein recently addressed a fundamental lack in the optogenetic toolbox and thereby opened up novel optoribogenetic applications^16^. Under blue light, the PAL receptor sequence-specifically binds to small RNA hairpins, denoted aptamers in the following, with up to around 100-fold enhanced affinity compared to darkness. Embedding these aptamers into choice RNAs enabled the optoribogenetic control of transcription initiation^17^, mRNA translation^16^, RNA interference^18^, and ribozyme activity^19^. More recently, the optoribogenetic concept was extended to controlling the RNA-binding activity of a homodimeric protein via coupling to a photoreceptor module that dimerizes under light^20^.

To facilitate the analysis and modulation of bacterial physiology, we here devised efficient setups for the optoribogenetic control of prokaryotic expression by the PAL receptor. Through optimization of promoters, translation initiation regions, and the PAL aptamer sequence, we generated the pCrepusculo setup which affords the downregulation of target gene expression under blue light. The pCrepusculo circuit is realized on one portable plasmid and employs the PAL receptor as a single polypeptide component. In contrast to existing systems for light-regulated gene expression in bacteria, PAL exerts optogenetic control at the mRNA level. Hence, PAL-based regulatory circuits lend themselves to the integration with circuitry, optogenetic or otherwise, acting at the DNA level. We demonstrated this concept by generating the derivative pAurora setup which inverts the system response at the DNA level and thus enables the strong upregulation of target genes under blue light. In a similar vein, the combination with a light-sensitive two-component system allowed expression to be bimodally activated under low blue light and deactivated again under strong blue light. Taken together, the optoribogenetic control at the mRNA level presented here expands the arsenal of optogenetics and thereby benefits diverse applications in microbial biotechnology and synthetic biology.

## Results and discussion

### A one-plasmid system for the optoribogenetic control of bacterial expression

We previously demonstrated that the light-activated, sequence-specific binding of PAL to its RNA aptamer can principally be harnessed to control bacterial expression at the mRNA level^16^. To this end, we interleaved the aptamer hairpin with the Shine-Dalgarno (SD) sequence, i.e. the ribosome-binding site, that governs the translation of an mRNA encoding a *Ds*Red fluorescent reporter (Fig. 1A). When combined with PAL, the reporter expression was diminished under blue light relative to darkness^16^. Although the previous setup enabled the efficient probing of PAL functional variants and showcased the basic validity of the optoribogenetic approach, it suffered from several shortcomings. First, the PAL receptor and the *Ds*Red reporter were encoded on two separate plasmids, thus requiring the use of two antibiotics. Second, PAL and *Ds*Red were expressed from the inducible pBAD and T7-*lacO* promoters, which are controlled by AraC and LacI, respectively. Addition of the chemical inducers L-arabinose and β-isopropyl-thiogalactoside (IPTG) was thus a necessity, and the system response strongly depended on the timing and extent of inducer addition. Moreover, given the use of the T7 promoter, the setup was only operational in bacterial cells expressing T7 RNA polymerase, e.g., via the DE3 lysogen. Third, the dynamic range of regulation, defined as the ratio of reporter gene expression under dark and blue-light conditions, was maximally 10-fold at a temperature of 29°C^16^ but strongly diminished to merely 1.8-fold at 37°C. Taken together, the earlier setup proved unwieldy and inefficient, thus complicating its practical application.

**Figure 1.**
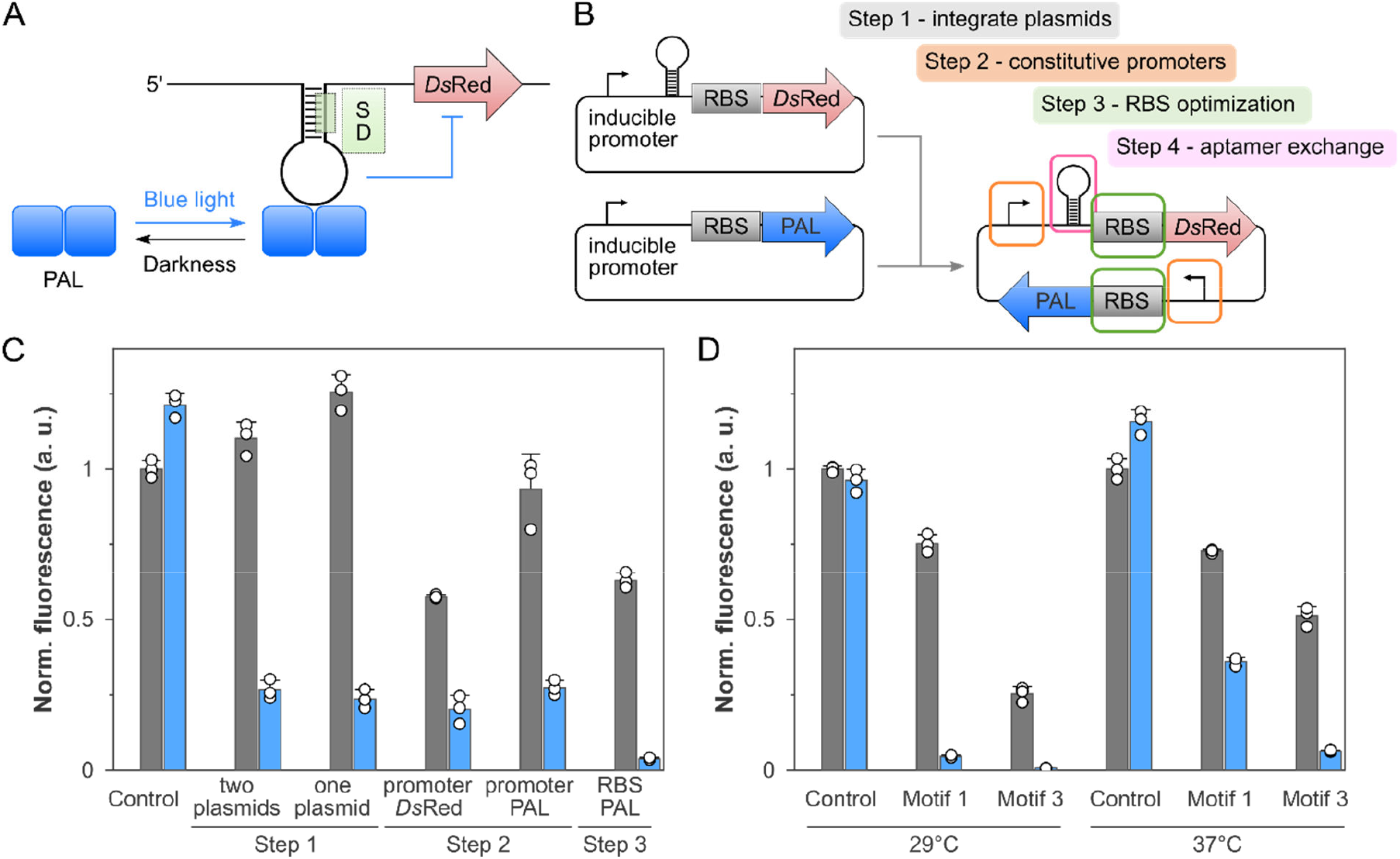
Implementation of one-plasmid systems for the optoribogenetic regulation of bacterial gene expression. **A**, Schematic of the system. The Shine-Dalgarno (SD) region upstream of a target gene, e.g., the red-fluorescent *Ds*Red, is interleaved with a small aptamer that forms a hairpin. Under blue light, the LOV receptor PAL binds to this aptamer and thus hinders translation. **B**, Strategy for the iterative optimization of bacterial optoribogenetic control. Two originally separate plasmids encoding the components of the system are combined onto a single backbone (step 1), followed by the optimization of promoters (step 2) and ribosome-binding sites (step 3) that govern PAL and target-gene expression. Finally, the original aptamer (motif 1) was replaced by a recently discovered variant (motif 3) with enhanced light-dependent interactions with PAL (step 4). **C**, To monitor and assess the iterative optimization, the different systems were transformed into *E. coli*, and reporter-gene expression was recorded for cultures grown at 29°C in either darkness (grey bars) or under blue light (blue bars). **D**, Plasmid variants with either the motif-1 or motif-3 aptamer were assessed at 29°C and 37°C. Data in panels C and D represent the mean ± s.d. of three biological replicates; a vector with omitted PAL was used as control. **E**, Fluorescence anisotropy of the binding of PAL to TAMRA-labeled motif 1 at 37°C in darkness (black dots) and under blue light (white dots). **F**, as in E but for TAMRA-labeled motif 3. Experiments in panels E and F were repeated twice with similar outcomes.

To overcome these deficits and to provide a more robust and accessible platform for the optoribogenetic control of bacterial expression, we iteratively optimized the setup as laid out in Fig. 1B. In a first step, we subcloned the *Ds*Red reporter cassette including the PAL aptamer onto the PAL expression plasmid, thereby assembling all system elements on a single plasmid backbone with a CDF origin of replication and a streptomycin resistance marker. To assess the effect of this and subsequent modifications, the respective plasmids were transformed into *E. coli* cells, and bacterial cultures were induced by addition of 1 mM IPTG and 4 mM L-arabinose, and were then grown at 29°C in either darkness or under 40 μW cm^-2^ blue light (470 nm) for 18 h. Doing so revealed that the integration of the system onto a single backbone entailed a slight increase of the dynamic range to 5.3 ± 0.7, compared to 4.1 ± 0.5 in the two-plasmid setup. We next replaced the inducible T7-lacO promoter of the *Ds*Red reporter cassette for a small library of constitutive promoters of varying strength^21,22^. Whereas introduction of the strong pH207 promoter abolished light regulation, systems with the promoters pJ5, pLλ, pSpc, and *rrnB* P1 retained light responses, albeit at somewhat reduced dynamic range of between 3- and 4-fold (Fig. 1C, Suppl. Fig. 1A). Based on the reporter expression in darkness, we selected the plasmid variants with the pSpc and pLλ promoters for further analysis. We next exchanged the inducible pBAD promoter that drives PAL expression in these plasmids for orthogonal constitutive promoters, which generally impaired the regulatory response. The best dynamic range of 3.4 ± 0.5 resulted for a combination of the pH207 promoter controlling PAL expression and the pLλ promoter governing *Ds*Red expression, and the following optimization was conducted with the corresponding plasmid.

Besides the promoter sequence, the ribosome-binding site (RBS) crucially determines the expression levels of target genes. We reasoned that modification of the RBS strengths of the PAL and *Ds*Red cassettes could prove a viable strategy for improving the dynamic range of light regulation. Given the relatively high basal *Ds*Red reporter expression under blue light, we suspected that an increase in PAL expression might lead to more efficient translational repression and thus entail an improved regulatory response. To this end, we used the RBS calculator server^23^ to evaluate and subsequently optimize the RBS sequences. In the case of the PAL expression cassette, the calculated translation initiation rate in the original design was 7,100 a.u. (arbitrary units). We then designed and tested RBS variants with predicted strengths of 20,000, 40,000, and 100,000 a.u. (Suppl. Fig. 1B). Consistent with the design rationale, all three RBS variants exhibited lower reporter fluorescence under blue light. In the variant with a predicted strength of 40,000 a.u., fluorescence was strongly suppressed in darkness as well. By contrast, the RBS strengths 20,000 a.u. and 100,000 a.u. largely preserved expression levels in darkness and hence gave rise to enhanced dynamic ranges of light regulation of 16.5- and 11.3-fold, respectively. We thus selected the former plasmid variant and next assessed the effect of altering the RBS that governs the expression of the target reporter gene. Notably, this RBS intentionally overlaps with the aptamer that PAL binds to under blue light, and hence the scope for its optimization is narrowly circumscribed. Using the RBS calculator, we designed RBS variants with predicted strengths of 2,500, 7,600, 13,600 a.u., compared to the calculated initial value of 5,700 a.u. for the original RBS. All these variants however displayed inferior regulation, with the first two variants abrogating light responsiveness altogether.

To further boost light-dependent regulation, we next considered a recently discovered aptamer variant, denoted motif 3^19^, which at 25°C exhibited more than 10-fold enhanced affinity for light-activated PAL than the formerly identified motif 1^16^ used in the experiments above. Introduction of the motif-3 aptamer consequently incurred reduced reporter fluorescence in darkness and, even more so, under blue light, thus resulting in an enhanced dynamic range of 33 ± 4 at 29°C (Figure 1D). We next assessed the performance of these systems at 37°C. For motif 1, the temperature increase was detrimental to the light response and decreased the dynamic range to a mere 2-fold, chiefly owing to impaired repression of reporter expression under blue light. Motif 3 also experienced elevated basal expression at 37°C compared to 29°C but retained a higher dynamic range of 8.0 ± 0.6. As this plasmid variant exhibited robust performance at both 29 and 37°C, we named it pCrepusculo. The variation between motifs 1 and 3 at the different temperatures could be principally due to differences in the stability and affinity of the RNA hairpins. To assess this concept, we next monitored the interaction between PAL and the RNA aptamers by fluorescence anisotropy at 37°C. To this end, the motif-1 and motif-3 aptamers were 5’-labelled with tetramethylrhodamine (TAMRA) and incubated with increasing concentrations of purified PAL protein in darkness or blue light. The rise in TAMRA fluorescence anisotropy resulting from PAL binding was fitted to single-site binding isotherms (Fig. 1E-F). As at 25°C, motif 3 exhibited tighter PAL interactions than motif 1 also at 37°C with *K*_d_ values of (270 ± 40) nM and (2.9 ± 0.5) nM in darkness and blue light, respectively. In comparison, the *K*_d_ values for motif 1 were (2600 ± 400) nM and (140 ± 20) nM. Notably, the affinity of PAL for motif 1 under blue light was thus around two-fold weaker at 37°C than at 25°C. By contrast, the affinity of light-activated PAL for motif 3 was essentially unaffected by the temperature increase. Taken together, the higher affinity of motif 3 and its less pronounced thermal sensitivity account for the superior performance of this aptamer in pCrepusculo.

### Inversion of the light response facilitates spatial control of gene expression

While light-repressed gene expression, as afforded by pCrepusculo, is principally suited for certain use cases, e.g., the optogenetic control of production processes in bacteria^24^, the application scope for light-induced expression appears yet larger. We thus set out to invert the blue-light response of pCrepusculo by inverter gene cassettes, as successfully implemented for previous optogenetic setups that control gene expression at the DNA level^25–27^. These inverters essentially encode a constitutively active bacterial transcription factor which strongly represses a downstream promoter, also included in the gene cassette. We tested three different inverters based on the λ phage cI^28^, SrpR, and PhlF repressors^29^. Upon insertion into pCrepusculo (Fig. 2A), the SrpR and PhlF inverters gave rise to elevated expression under blue light at 29°C but failed to much repress expression in darkness (Fig. 2B). Consequently, the dynamic range of light regulation (i.e. expression under blue light vs. darkness) was less than two-fold. At 37°C, the SrpR and PhlF inverters fared even worse, and constitutively high expression resulted with no significant light response observable. By contrast, the λ cI inverter achieved robust up-regulation of expression under blue light at both 29°C and 37°C, with dynamic ranges of 16 ± 2 and 18 ± 6, respectively. We hence continued the experiments with the corresponding plasmid and dubbed it pAurora.

**Figure 2.**
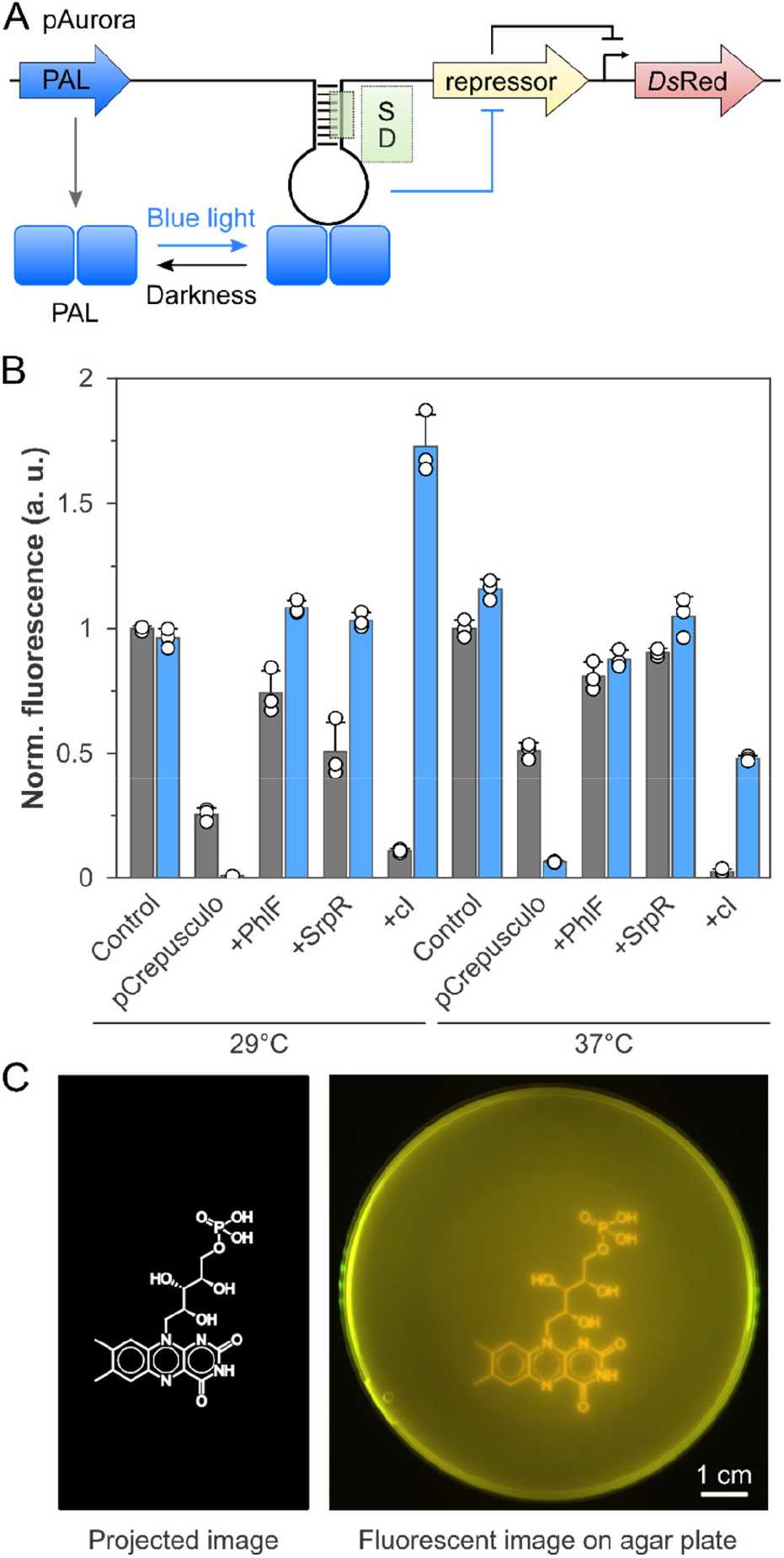
Construction of the pAurora system with inverted light response. **A**, Schematic of the pAurora system. In contrast to pCrepusculo, *Ds*Red expression is under the control of a repressor protein, the expression of which in turn is regulated by PAL. Under light, PAL suppresses the repressor expression and thereby prompts *Ds*Red expression. **B**, Different constructs en route to the eventual pAurora system were evaluated for light-dependent reporter expression at 29°C and 37°C, with cultures grown in darkness and under blue light shown as grey and blue bars, respectively. Three different inverter cassettes based on the SrpR, PhlF, and λ cI repressors were assessed. Data represent mean ± s.d. of three biological replicates, and the control refers to a plasmid variant with PAL omitted. **C**, Bacteria carrying the pAurora plasmid were embedded in agar and illuminated overnight with a photomask (left). Afterwards, the expression of the *Ds*Red reporter inside the bacteria was visualized by fluorescence (right). The scale bar denotes a distance of 1 cm.

We next investigated whether pAurora supports the spatially resolved control of gene expression. As pioneered by Voigt and colleagues^30^, optogenetically regulated gene expression can not least be harnessed for so-called ‘bacterial photography’ in which an image is projected onto a bacterial lawn. To this end, we embedded bacteria harboring pAurora into agar and incubated them overnight at 37°C under constant blue light (470 nm, 5 μW cm^-2^). During incubation, the agar plate was covered by a transparency displaying the Lewis formula of flavin mononucleotide, the chromophore of PAL and other LOV receptors. Fluorescence analysis after incubation showed that expression of the *Ds*Red reporter was spatially confined to the illuminated areas, and thus the projected image was faithfully reproduced by the bacterial lawn. Despite the simplicity of the illumination setup, spatial features down to approximately 300 μm were accurately resolved.

### Quantitative analysis of optoribogenetically controlled bacterial expression

To glean additional insight into the efficiency and sensitivity of light-dependent gene expression, we analyzed the pCrepusculo and pAurora systems in more detail. First, we determined light dose-response curves by incubating bacteria harboring one of the two systems under different blue-light intensities at 37°C (Fig. 3A). In case of pCrepusculo, the reporter gene expression decreased monotonically with light intensity by a factor of around 8-fold. A fit of the data to a hyperbolic function yielded a half-maximal light dose *I*_50_ of (0.6 ± 0.2) μW cm^-2^. For pAurora, reporter expression hyperbolically increased with blue light by almost 70-fold with a half-maximal dose of (1.2 ± 0.1) μW cm^-2^. Second, we probed how rapidly pCrepusculo and pAurora respond to changes in illumination. To this end, bacteria containing either plasmid were first cultivated under non-inducing conditions, i.e. 40 μW cm^-2^ blue light for pCrepusculo and darkness for pAurora. At an optical density at 600 nm of around 0.4, the cultures were transferred to inducing conditions, i.e. darkness for pCrepusculo and 40 μW cm^-2^ blue light for pAurora. Samples, taken at this point and at different times afterwards, were arrested by antibiotic addition and analyzed for *Ds*Red fluorescence (Fig. 3B). In both systems, the reporter expression increased sigmoidally with the time spent at inducing conditions. Fits to logistic functions yielded half-maximal activation times *t*_50_ of (2.3 ± 0.4) h and (2.5 ± 0.2) h for pCrepusculo and pAurora, respectively. Notably, the sigmoidal time course was shallower for pCrepusculo, with an onset of the fluorescence increase already apparent at 30 minutes compared to around 90 minutes for pAurora. It is worth noting that different biological processes underlie the lag phase until gene expression ramps up in the two systems. In pCrepusculo, the rate-limiting step is the slow recovery of PAL from its light-adapted state to the dark-adapted state, which at 37°C proceeds with a time constant of around 400 s (Suppl. Fig. 2). By contrast, in pAurora the reporter induction is governed by several steps including the depletion of the λ cI repressor and the onset of transcription of the reporter. These considerations may account for the different induction time courses in pCrepusculo and pAurora.

**Figure 3.**
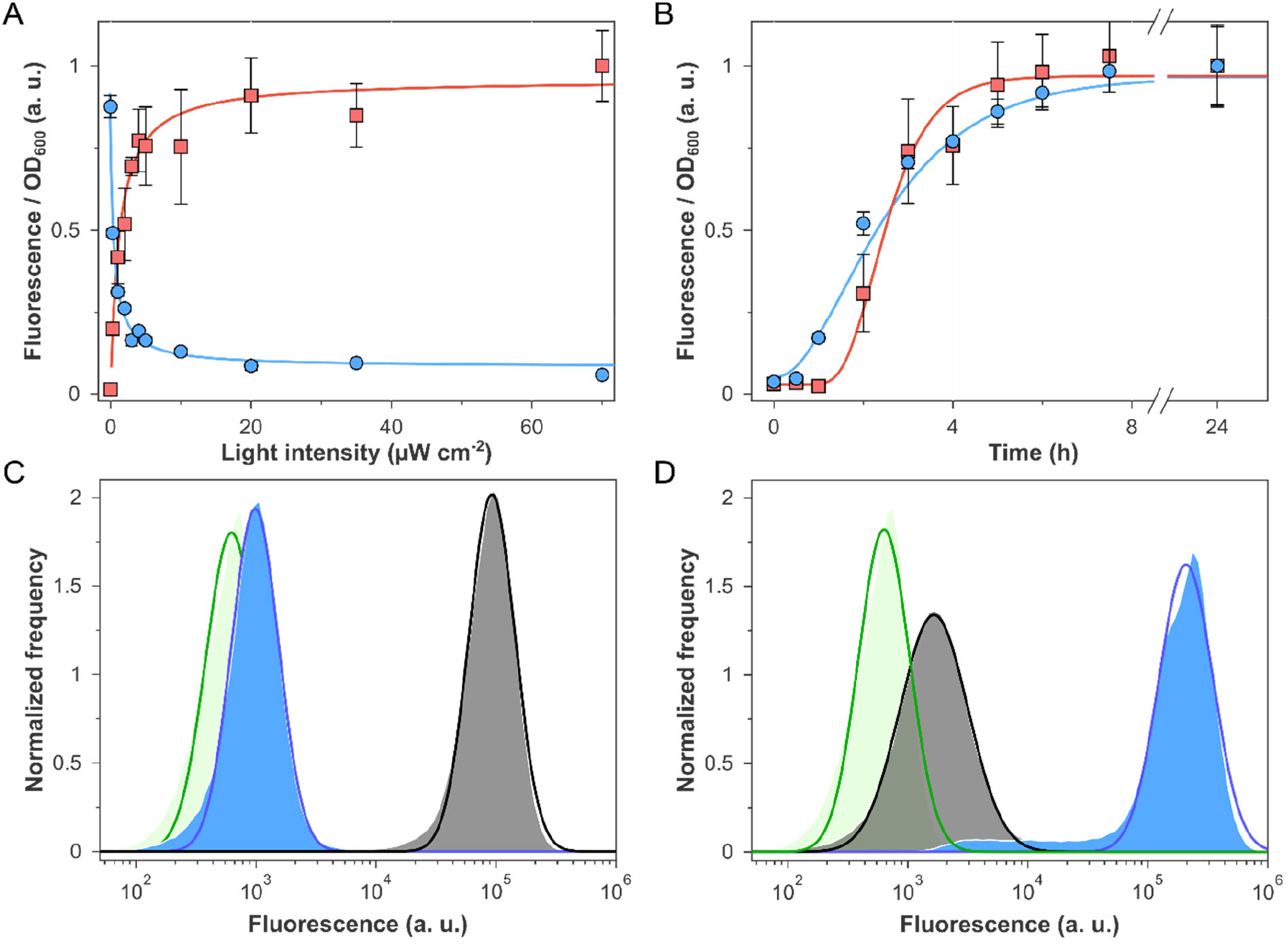
Characterization of the pCrepusculo and pAurora systems. **A**, Bacterial cultures harboring the pCrepusculo (blue symbols and curves) and pAurora plasmids (red) were grown under different blue-light intensities at 37°C for 24 hours, followed by determination of the resultant *Ds*Red fluorescence. **B**, Bacterial cultures containing pCrepusculo and pAurora were grown under non-inducing conditions until an optical density at 600 nm of 0.4 was reached. The cultures were then transferred to inducing conditions, i.e. darkness for pCrepusculo and blue light for pAurora, and samples were drawn and arrested by antibiotic addition at different times. After allowing for *Ds*Red maturation, the fluorescence was recorded. Data in panels A and B represent mean ± s.d. of three biological replicates. **C**, Flow cytometry analysis of bacterial cultures harboring pCrepusculo and cultivated for 24 h at 37°C in darkness (grey curve) or under 60 μW cm^-2^ 470-nm light (blue). The green curve represents bacteria carrying an empty vector. **D**, As in C but for pAurora. The experiments in C and D were repeated three times with similar results.

The above fluorescence analyses were all performed at the culture level, thus reporting on an ensemble average of *Ds*Red production, without information on the homogeneity of the samples. We hence evaluated the light responses of bacteria harboring pCrepusculo and pAurora at the single-cell level by flow cytometry (Fig. 3C,D). In case of pCrespuculo, we observed homogenous log-normal distributions with median fluorescence values of 10^5.0^ a.u. and 10^3.0^ a.u. for bacteria cultivated in darkness and under 60 μW cm^-2^ blue light, respectively, corresponding to a 94-fold change. Notably, the basal expression of pCrepusculo in darkness was only slightly elevated compared to an empty-vector control with median fluorescence at 10^2.8^ a.u. For pAurora, the median fluorescence was 10^3.2^ a.u. for bacteria grown in darkness and 10^5.3^ a.u. for those grown under blue light, which corresponds to a 126-fold difference. The measurements on light-adapted pAurora identified a minor population with lower single-cell fluorescence which may reflect partial plasmid loss or incapacitation during growth. Nonetheless, the flow-cytometry data indicate that both pCrepusculo and pAurora respond to illumination stringently and largely in all-or-none fashion.

### Parallel optogenetic control at the DNA and mRNA levels

The pCrepusculo and pAurora systems achieve light control at the mRNA level, as opposed to other setups for light-controlled bacterial expression which generally operate at the DNA level. We reasoned that, owing to this principal difference, pCrepusculo and pAurora may particularly lend themselves for combinations with other optogenetic systems. To evaluate this concept, we cloned the YPet yellow-fluorescent reporter into our plasmids, yielding pCrepusculo-YPet and pAurora-YPet. We then cotransformed these plasmids with the pDusk and pDawn plasmids^25^ which underlie manifold optogenetic applications in bacteria^24,31–37^ (Fig. 4A). Briefly, pDusk encodes the blue-light-sensitive two-component system (TCS) YF1:FixJ^38^ and mediates downregulation of a reporter gene, here *Ds*Red, under blue light compared to darkness. The pDawn-*Ds*Red plasmid derives from pDusk-*Ds*Red and inverts and amplifies the light response via a λ-cI-based inversion cassette^25^. Bacteria carrying either pCrepusculo-YPet or pAurora-YPet combined with either pDusk-*Ds*Red or pDawn-*Ds*Red were incubated at 37°C in darkness or under 60 μW cm^-2^ blue light, followed by determination of the YPet and *Ds*Red reporter fluorescence. By and large, the individual optogenetic systems exhibited similar light responses when combined with either another optogenetic system or an empty-vector control. As an exception, the dynamic range of light regulation was suppressed for pCrepusculo-YPet when combined with pDawn-*Ds*Red which we attribute to the presence of the strong λ cI repressor encoded on pDawn. Notably, both the pAurora and pDawn plasmids use the same inverter to alter the response to light, and crosstalk between them is hence expected. That notwithstanding, in combination these plasmids mediated strong induction of both the YPet and *Ds*Red reporter genes under blue light. Taken together, these data indicate that pCrepusculo and pAurora can be deployed in combination with at least certain optogenetic setups that achieve regulation solely at the DNA level.

**Figure 4.**
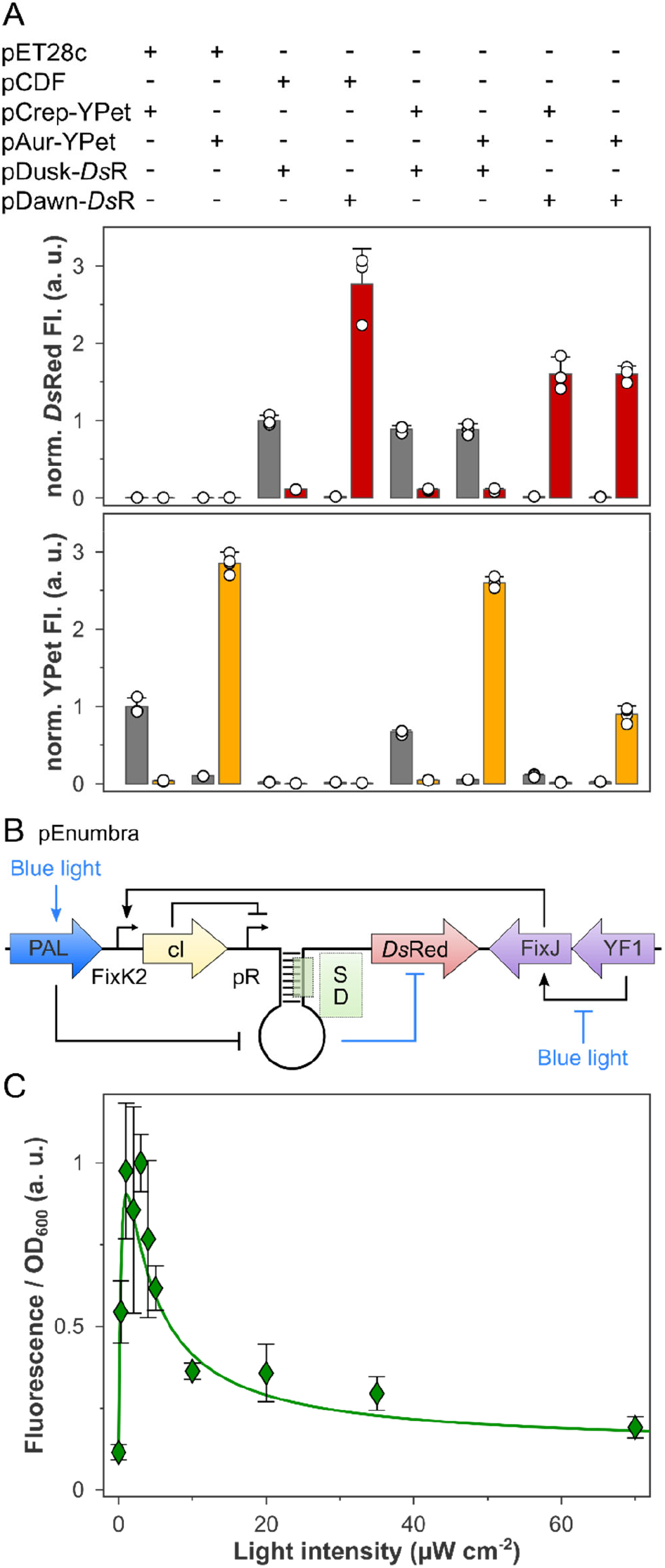
Combined optogenetic regulation at the DNA and mRNA levels. **A**, Bacterial cells were transformed with different combinations of pCrepusculo and pAurora with the pDusk and pDawn plasmids. In these experiments, the pCrepusculo and pAurora plasmids drive the expression of the yellow-fluorescent reporter YPet, whereas pDusk and pDawn use the red-fluorescent *Ds*Red. pET28c and pCDF denote empty-vector controls. Bacterial cultures were grown at 37°C in darkness or under blue light (470 nm, 60 μW cm^-2^), and the fluorescence of *Ds*Red (upper panel) and YPet was measured (lower panel). **B**, Schematic of the pEnumbra plasmid that integrates the PAL and YF1:FixJ regulatory circuits. **C**, Bacteria containing pEnumbra were grown at 37°C under different light intensities for 24 h, and *Ds*Red reporter fluorescence was determined. Data in panels A and C represent mean ± s.d. of three biological replicates.

Given that PAL acts at the mRNA level, we next sought to devise circuits that integrate translational and transcriptional optogenetic regulation. Such circuits might exhibit altered responses to light and offer finer-grained control. To this end, we placed the PAL expression cassette onto the pDusk and pDawn backbones and embedded the PAL aptamer into the Shine-Dalgarno sequences controlling *Ds*Red expression in these plasmids. We hypothesized that doing so might achieve dual regulation of expression by the YF1:FixJ TCS on the one hand and PAL on the other hand (Fig. 4B). We also constructed separate plasmid variants that employed motif 1 rather than the motif-3 aptamer. We next recorded reporter fluorescence for cultures harboring these plasmids upon cultivation in darkness or blue light at intensities of 5 and 60 μW cm^-2^ (Suppl. Fig. 3). Within the pDusk context, introduction of the aptamers reduced *Ds*Red expression under all light conditions. Whereas the plasmid variant with the motif-1 aptamer had a similar dynamic range as the parental pDusk, for motif 3 an almost two-fold improvement was achieved, chiefly owing to pronounced repression under blue light. In case of the pDawn plasmid variants, the reporter gene is antagonistically regulated, with activation of the YF1:FixJ TCS resulting in higher expression, and PAL activation prompting lower expression. At a light intensity of 5 μW cm^-2^, the reporter fluorescence increased by at least 8-fold compared to darkness for both motifs. By contrast, more intense blue light (60 μW cm^-2^) resulted in an up to 4-fold decrease of fluorescence. These data likely reflect that the YF1:FixJ TCS is triggered at low light intensities before the PAL receptor is fully activated at higher light intensities. We hence further derivatized the plasmids by exchanging the residue Val28 in YF1 for Ile^39,40^ which engenders a tenfold slowdown of dark recovery and a concomitant increase of effective light sensitivity at photostationary state^40,41^. Doing so led to a slight increase of reporter fluorescence at 5 μW cm^-2^ blue light while the values in darkness and at 60 μW cm^-2^ were little affected.

Given the performance of the individual systems, we selected the pDawn-*Ds*Red plasmid with the motif-3 aptamer and the V28I substitution for further analysis and named it pEnumbra. To probe the properties of this system in more detail, we recorded a light dose-response curve for bacteria harboring pEnumbra (Fig. 4C). In darkness, *Ds*Red reporter fluorescence was low but increased by up to around 10-fold at light intensities between 1 and 2 μW cm^-2^, which we attribute to activation of the highly sensitive V28I variant of YF1 even at low light levels. At higher light intensities, the expression dropped by around 5-fold compared to its maximal value. This response can be explained by subsequent activation of the PAL receptor and repression at the mRNA level. Taken together, the pEnumbra system thus achieves bimodal expression control with but a single light color.

## Conclusion

Although a plethora of photoreceptors now exist that permit the optogenetic control of diverse cellular aspects and processes, light-regulated gene expression remains particularly versatile and widely applied in both prokaryotes and eukaryotes^42–44^. Prominent use cases for light-regulated gene expression in bacteria include the modulation of production processes^24,44^, the spatial patterning of expression and the templating of materials^36,37^, and the control of bacteria inside the digestive tract of animals^35,45^. The dominant strategies for achieving optogenetic control of bacterial gene expression employ either light-sensitive two-component systems^25–27,30,46^, protein-protein interactions^47–49^, or second-messenger signaling^50^. All these setups have in common that they operate at the DNA level and involve the regulation of transcription initiation. The LOV receptor PAL^16^ stands apart in that it acts at the RNA level. Through iterative optimization, we have devised the pCrepusculo setup in which PAL associates with a small RNA hairpin near the ribosome-binding site of target genes and thus represses their expression under blue light by around 8-to 30-fold depending on temperature. While the dynamic range is less than in certain of the transcriptional optogenetic approaches^25,27,46,47^, it compares to or exceeds the regulatory effect realized for the control of bacterial translation initiation by ligand-responsive riboswitches, e.g., references^51–53^.

The light-dependent expression control at the mRNA level offers at least two principal advantages that can be leveraged for applications in optogenetics, biotechnology, and synthetic biology. First, as regulation occurs downstream of transcription, responses to illumination are expected to manifest faster than for optogenetic approaches acting at the DNA level. This notion is supported by the comparatively fast onset of induction evidenced for pCrepusculo upon withdrawal of blue light (see Fig. 3B). Second, PAL-based regulation is orthogonal to mechanisms, be they optogenetic or not, which target transcription, particularly its initiation. Therefore, the optoribogenetic control afforded by PAL can readily be combined and integrated with DNA-based regulatory approaches. In one demonstration of this concept, we have devised the pAurora plasmid which inverts and amplifies the light response of pCrepusculo via an inverter gene cassette. The ability to activate, rather than depress, expression by light benefits multiple applications as showcased for the spatial control of protein production (see Fig. 2C). Moreover, we embedded the PAL-based optoribogenetic system into the widely used pDusk and pDawn plasmids and thus established responses to blue light that could not be obtained with either circuit alone. In specific, the pEnumbra setup was bimodally controlled, with strong expression at intermediate illumination levels but lower expression in darkness and under more intense light. Put another way, pEnumbra thus constitutes a sensor circuit that responds to discrete light-intensity bands. The bimodal control by a single light color will be especially useful for multiplexed applications of optogenetic setups and other light-responsive elements, e.g., fluorescent proteins. More broadly, the combination with PAL principally extends to scores of other systems for the optogenetic control of gene expression and could thus install an added layer of control. The implementation and characterization of a new, compact RNA aptamer (motif 3) that interacts with PAL in strongly light-regulated fashion paves the way towards other modes of regulation at the RNA level as already well established for riboswitches^54^. Notably, the PAL:motif-3 interaction surpasses a recently developed light-responsive RNA:protein pair^20^ in terms of both overall affinity and regulatory response to light. Therefore, the motif-3 aptamer stands to also benefit optoribogenetics in mammalian cells^16–18^. In addition to deploying PAL unmodified, the application scope can be further expanded by covalent attachment of diverse effector modules, e.g., intracellular localization tags or RNA-modifying enzymes. As detailed in the original report on PAL^16^, it is imperative that any such modules are connected to the N terminus of PAL because the C terminus is centrally involved in light-dependent signal transduction. Covalent linkage of modules to the PAL C terminus, as recently performed^20^, greatly impairs light responsiveness.

## Methods

### Molecular biology

The plasmids pCDF-PALopt and pET-28c-*Ds*Red-SP^16^ were the starting points for the construction of pCrepusculo. First, the expression cassette encoding the *Ds*Red Express 2 reporter gene^55^ was subcloned into the pCDF-PALopt vector. This and all subsequent cloning steps were carried out by Gibson assembly^56^, unless remarked otherwise. The inducible promoters controlling expression of PAL and *Ds*Red, respectively, were next replaced by constitutive ones (rrnB P1^21^, PJ5, PL Lambda, PSpc, and PH207^22^). Then, the plasmid with the PH207 and PL Lambda promoters controlling PAL and *Ds*Red, respectively, had the ribosome-binding sequences (RBS) altered. Finally, the aptamer sequence originally used in the pET-28c-*Ds*Red-SP plasmid (motif 1) was exchanged for that of motif 3 to yield pCrepusculo. For the construction of pAurora, an inverter gene cassette based on the λ phage cI repressor was amplified by PCR from the pDawn plasmid^25^ and inserted into pCrepusculo. Doing so put cI expression under the control of PAL, while the *Ds*Red gene was placed downstream of the pR promoter that is controlled by the repressor. Inverter cassettes based on the SrpR and PhlF repressors were obtained as synthetic genes (GeneArt, ThermoFisher) and were likewise introduced into pCrepusculo. Empty versions of pCrepusculo and pAurora in which a multiple cloning site (MCS) replaces the fluorescent reporter gene were generated by PCR amplification and blunt-end ligation and deposited with Addgene under accession numbers XXX and YYY.

To assess the combination with pDusk and pDawn, a synthetic gene encoding the YPet fluorescent reporter was ordered with *Escherichia coli*-adapted codon usage (GeneArt) and cloned into pCrepusculo and pAurora. For the construction of pEnumbra and related plasmids, PCR amplification and blunt-end ligation were used to insert the motif-1 or motif-3 aptamers into the pDusk and pDawn plasmids near the ribosome-binding sites controlling the *Ds*Red reporter. The PAL expression cassette was then amplified by PCR from pCrepusculo and inserted into the plasmids. To introduce the V28I substitution, a corresponding pDawn variant^40^ was used as the starting plasmid.

For protein expression, the PAL gene with codons optimized for *E. coli* was subcloned from the pET-28c-PALopt plasmid^16^ into a pET-19b backbone. In the resultant plasmid pET-19b-SUMO-PAL, the gene was thereby equipped with an N-terminal hexahistidine sequence followed by a SUMO tag (small ubiquitin-like modifier). The identity of all constructs was confirmed by Sanger DNA sequencing (Microsynth, Göttingen).

### Bacterial reporter assay

The reporter assays for the original two-plasmid system and subsequent iteratively optimized setups were performed in 14-mL culture tubes. *E. coli* CmpX13^57^ cells harboring the desired plasmid constructs were inoculated in 5 mL lysogeny broth (LB) plus the respective antibiotic, i.e. 50 μg mL^-1^ kanamycin and 100 μg mL^-1^ streptomycin. Bacterial cultures were grown overnight under non-inducing conditions, i.e., depending on the construct, either in darkness or under 40 μW cm^-2^ 470-nm blue light. Following incubation, 50 μL of each culture were used to inoculate two fresh 5-mL liquid cultures, with 4 mM L-arabinose and 1 mM β-isopropyl thiogalactoside (IPTG) added as inducers if necessary. The samples were then shaken at 225 rpm in either darkness or under 40 μW cm^-2^ 470-nm light for 18 h and at 29°C or 37°C, depending on the performed assay. For assessing the motif exchanges, the introduction of inverter gene cassettes, the light dose response, and combinations with the pDusk and pDawn systems, bacteria were cultivated in deep-well 96-well blocks rather than in culture tubes. Each well contained 500 μL LB medium with the corresponding antibiotic, and following inoculation, the plate was incubated overnight at 37°C under non-inducing conditions with shaking set to 600 rpm. 198 μL LB medium plus the respective antibiotic were then mixed with 2 μL of the overnight starter culture and transferred to a 96-well plate. The plates were incubated for 24 h at 29°C or 37°C in darkness or under blue light at different intensities while shaking at 800 rpm. For both assay formats, the fluorescence of *Ds*Red [excitation at (554 ± 9) nm and emission at (591 ± 20) nm] and YPet [excitation at (500 ± 9) nm and emission at (530 ± 20) nm] was measured with a Tecan M200 multimode microplate reader and normalized by the optical density at 600 nm (*OD*_600_). All measurements are reported as mean ± s.d. of at least three biologically independent samples.

### Light-dose response curves

The assay was performed in 96-well deep-well microtiter plates at 37 °C as described above. The plates were illuminated with blue light (470 nm) at intensities of 0.3, 1, 2, 3, 4, 5, 10, 20, 35, and 70 μW cm^-2^. For pCrepusculo and p Aurora, the data were fitted to a hyperbolic function (eq. 1) with the Fit-o-mat software^58^.

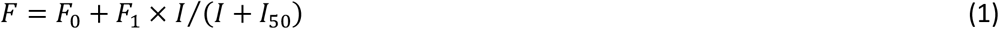

where *F* is the fluorescence value, *I* signifies the light intensity, and *I*_50_ is the light intensity which produces a half-maximal fluorescence signal.

In the case of pEnumbra, the data were fitted to a more complex model that considers light-induced activation of expression by the YF1:FixJ TCS circuit and its light-induced repression by the PAL circuit. Each circuit is assumed to react to light separately, i.e. with no cooperativity between them. The resultant relationship corresponds to the random-order mechanism of a two-substrate enzyme according to eq. 2:

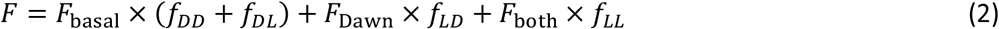

where *F*_basal_, *F*_Dawn_, and *F*_both_ are the fluorescence outputs if neither circuit, just the YF1:FixJ TCS circuit alone, or both circuits are activated by light together, respectively. The fractional populations *f* of the different activity states of the integrated circuit are given by eqs. (3-6).

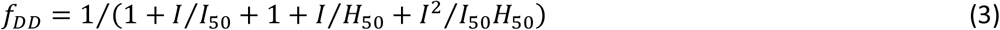

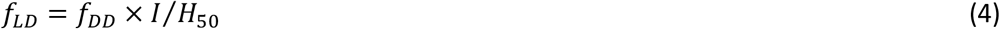

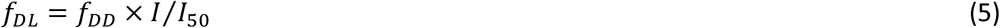

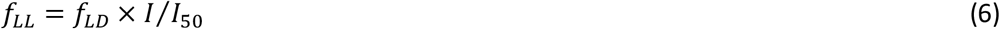

where the first letter in the subscript denotes the activity status of the YF1:FixJ TCS, i.e. D for the dark-adapted, basal state, and L for the light-adapted state. The second letter in the subscript denotes the corresponding activity status for the PAL circuit. The half-maximal light intensities *H*_50_ and *I*_50_ signify the light intensities *I* at which the YF1:FixJ and PAL circuits, respectively, are activated to 50%. When fitting the data, the parameter *I*_50_ was fixed at the value determined for the isolated PAL circuit within the pCrepusculo system.

### Induction kinetics

To record the induction kinetics of pCrepusculo and pAurora, triplicate starter cultures of each construct were grown overnight in 5 mL LB medium, supplemented with 100 μg mL^-1^ streptomycin, under non-inducing conditions (either darkness or 40 μW cm^-2^ 470-nm light) at 37°C and 225 rpm shaking. Triplicate 100-mL LB/Strep cultures were inoculated with the starter cultures and grown to an *OD*_600_ of ~ 0.4 under non-inducing conditions. Then, the cultures were transferred to inducing conditions, and samples were taken after 0, 0.5, 1, 2, 3, 4, 5, 6, 8, and 24 h. Cell growth and protein expression was inhibited immediately after sampling by addition of 0.4 mg mL^-1^ tetracycline and 3.5 mg mL^-1^ chloramphenicol^25^. The samples were stored on ice for at least 2 h to allow for *Ds*Red maturation^55^, followed by fluorescence measurements, as described above. The normalized fluorescence was fitted to a logistic function^25^ (eq. 7) to determine the time *t50* at which the half-maximal gene expression is reached.

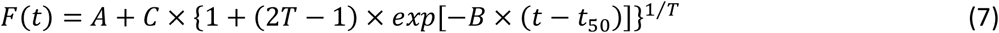

In eq. (7), the parameters *A, B, C*, and *T* determine the initial and starting values of the logistic function and its steepness.

### Bacterial photography

CmpX13 bacteria harboring the pAurora-*Ds*Red plasmid were grown overnight on an LB/Strep agar plate at 37°C. A single bacterial colony was used to inoculate a 5-mL LB/Strep culture, followed by incubation at 37°C for 4 hours in darkness. The culture was then rapidly mixed with 50 mL of molten LB-agar supplemented with 100 μg mL^-1^ streptomycin and poured into a Petri dish. The solidified agar plate was incubated for 2 h at 37°C in darkness and then transferred to a 3D-printed housing with an opening on top for a laser-printed transparency of the desired image. The entire contraption was incubated for 16 hours at 37°C with constant illumination of 5 μW cm^-2^ at 470 nm. The fluorescence of the bacteria on the agar plate was then imaged using a transilluminator FG-09 (Nippon Genetics), equipped with LED excitation at 470 nm and a long-pass emission filter at 520 nm.

### Flow cytometry

Triplicate cultures of pCrepusculo, pAurora, and the pCDF empty-vector control were grown in microtiter plates in darkness or under constant blue light (470 nm, 60 μW cm^-2^) for 24 h. Cells were then diluted in phosphate-buffered saline (1x sheath fluid, BioRad) and analyzed on a S3e Cell sorter (BioRad) based on their *Ds*Red fluorescence [excitation at 488 nm and 561 nm, emission (585 ± 15) nm]. Single-cell fluorescence distributions were fitted to log-normal functions.

### Expression and purification of PAL

The PAL protein was heterologously expressed in *E. coli* and purified by immobilized metal ion chromatography. Briefly, *E. coli* CmpX13 cells^57^ harboring the pET-19b-SUMO-PAL expression plasmid (see above) were grown at 37°C in lysogeny broth medium supplemented with 50 mg mL^-1^ ampicillin and 50 μM riboflavin. Once an optical density at 600 nm of around 0.6 was reached, the temperature was lowered to 18°C and incubation continued for 1 h. Next, expression was induced by addition of 0.2 mM isopropyl *β*-D-1-thiogalactopyranoside. Following 16 h of incubation at 18°C, the cells were harvested by centrifugation, resuspended in buffer [50 mM Tris-HCl pH 8.0, 1 M NaCl, 10 mM imidazole, 1 mg mL^-1^ lysozyme, protease inhibitor cocktail (Roche)], and lysed by sonication. The cleared lysate was subjected to Co^2+^ immobilized metal ion affinity chromatography (IMAC). Bound protein was eluted with an imidazole gradient and pooled. The His-tagged SUMO protease Senp2 was added, and the sample was dialyzed against 10 mM Tris-HCl pH 8.0, 150 mM NaCl, 5% glycerol. Following dialysis, the PAL protein was purified using a second IMAC step and concentrated by spin filtration. The protein concentration was determined by UV/vis absorbance spectroscopy on an Agilent 8453 diode-array spectrophotometer using an extinction coefficient of 12,500 M^-1^ cm^-1^ at 450 nm.

### UV/vis spectroscopy and fluorescence anisotropy

Measurements of the recovery reaction of PAL after light exposure were conducted in 12 mM HEPES-HCl pH 7.7, 135 mM KCl, 10 mM NaCl, 1 mM MgCl2, 10% (v/v) glycerol, 100 μg mL^-1^ BSA^16^. PAL was first exposed to saturating blue light, and then its return to the dark-adapted state was monitored by acquiring serial spectra over time using an Agilent 8453 spectrophotometer. The absorbance values at 450 nm were corrected by those at 600 nm, and the resultant kinetics were fitted by single-exponential functions. Due to slow precipitation of PAL at elevated temperatures, precise measurements at 37°C were precluded. Hence, the rate constant for dark recovery *k*_-1_ was determined at 18°C, 22°C, 26°C, and 30°C. The dark recovery rate constant was then extrapolated to 37°C by fitting the experimental data to the Arrhenius equation (8) using the Fit-o-mat program^58^.

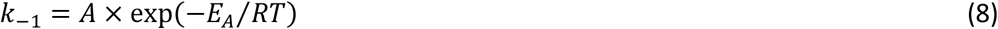

where *A* is the pre-exponential factor and *E_a_* the activation energy. The 95% confidence interval of the extrapolated curve was determined by bootstrapping^58^.

To probe RNA affinity, motif-1 (5’-ACCUUGGUUGAAGCAGACGACC-3’) and motif-3 (5’-ACCUUCGGUUCAGCAGCGAGCCG-3’) aptamers were synthesized with a 5’-attached tetramethylrhodamine (TAMRA) fluorophore (Integrated DNA Technologies). 4 nM of the fluorescently labelled aptamer were incubated with increasing amounts of PAL in its dark-adapted state, and TAMRA fluorescence anisotropy was recorded at 37°C in black 384-well microtiter plates. Then, the samples were exposed to saturating blue light (470 nm, 10 mW cm^-2^) for one minute, and fluorescence anisotropy was recorded again. Fluorescence measurements were conducted with a CLARIOSTAR multimode microplate reader using (540 ± 10) nm excitation and (590 ± 10) nm emission filters, and a 566-nm long-pass filter. Anisotropy data were fitted to binding isotherms according to eq. (9).

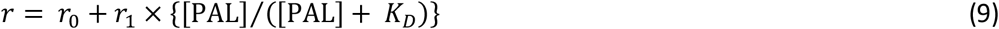

where *r*, [PAL], and *K*_D_ denote the fluorescence anisotropy, the PAL concentration, and the dissociation constant, respectively. In case of motif 3, the dissociation constant under blue light was in the same range as the PAL concentrations used. Therefore, a modified single-site model was employed to account for the depletion of free PAL (eq. 10)

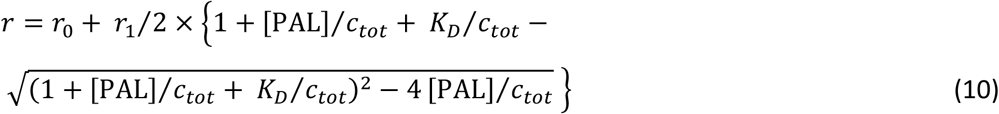

where *C*_tot_ is the total labelled-RNA concentration in the assay.

## Acknowledgements

Financial support by the Deutsche Forschungsgemeinschaft (grants MA3442/5-1 and MA3442/5-2 to G. M., and MO2192/6-1 MO2192/6-2 to A.M.) and by the European Union (ERC ‘OptoRibo’, 615381 to G.M.) is gratefully acknowledged. The pCrepusculo-MCS and pAurora-MCS plasmids will be available at Addgene under accession numbers XXX and YYY.

## Supplementary Information

**Suppl. Figure 1.**
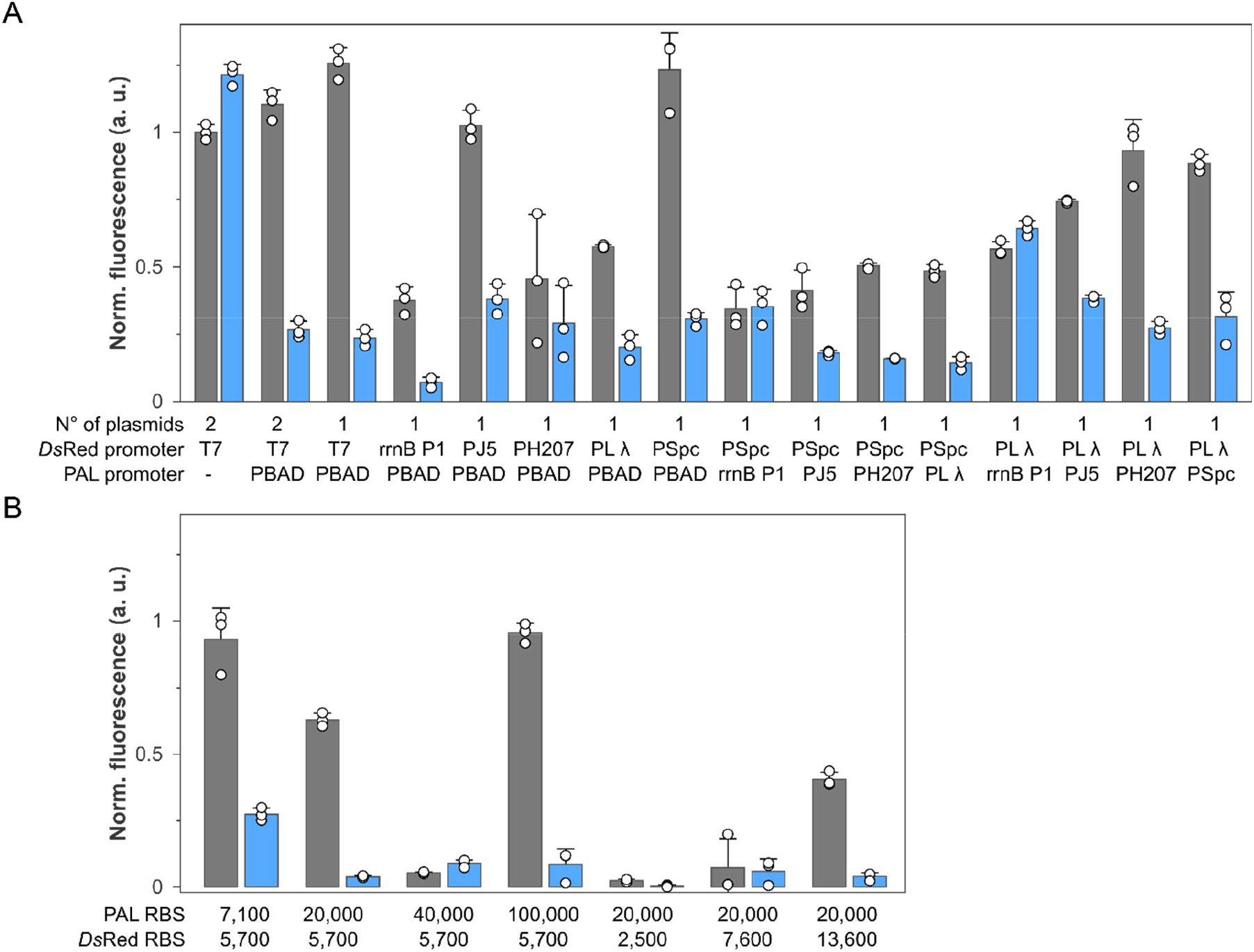
Iterative optimization of the optoribogenetic control of bacterial expression. **A**, Plasmid systems from different optimization steps were transformed into *E. coli* CmpX13, and reporter-gene expression was recorded for cultures grown at 29°C in either darkness (grey bars) or under blue light (blue bars). The number of different plasmids and the promoters driving the expression of PAL and the *Ds*Red reporter in the individual systems, respectively, are marked below the graph. **B**, Single-plasmid variants in which the ribosome-binding sites (RBS) determining PAL and *Ds*Red expression were adjusted. The predicted strength of the RBS is noted below the graph in arbitrary units (a.u.). Data in panels A and B represent the mean ± s.d. of three biological replicates; a vector with omitted PAL was used as control.

**Suppl. Figure 2.**
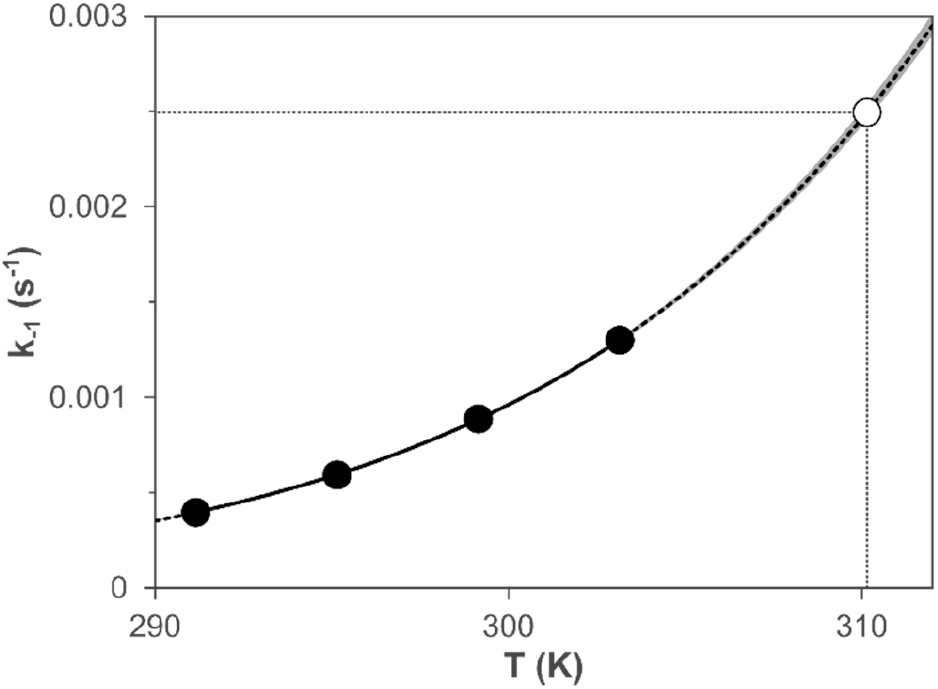
Analysis of the PAL dark-recovery reaction. The recovery rate constants *k*_-1_ determined at different temperatures between 14°C and 30°C (black dots) were fitted to the Arrhenius equation (solid line). The activation energy amounted to (72.8 ± 0.3) kJ mol^-1^, and extrapolation (dashed line, with 95% confidence interval indicated by grey shading) yielded a recovery rate constant of 0.0025 s^-1^ at 37°C (white dot). Data represent the mean of two independent measurements.

**Suppl. Figure 3.**
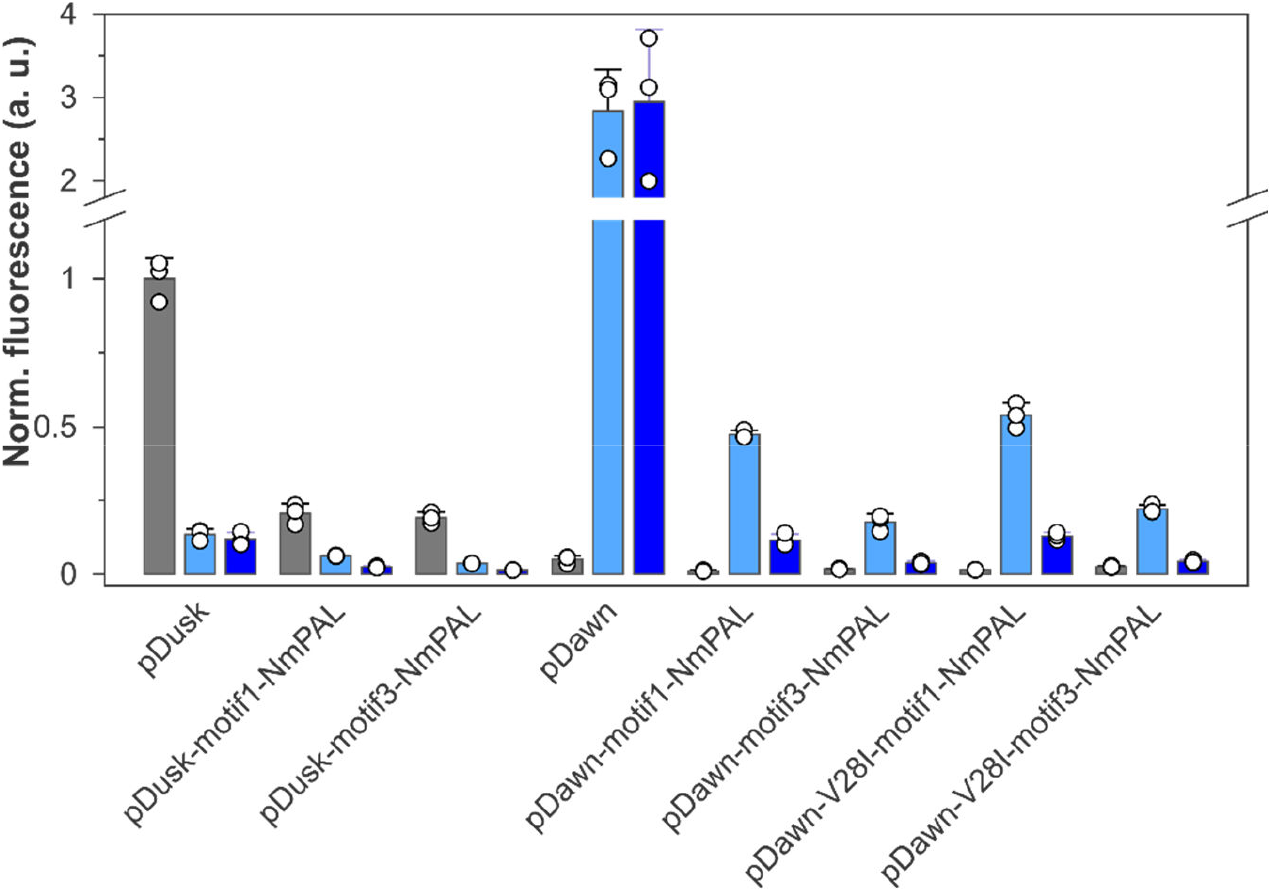
Combined optogenetic control with PAL and the YF1:FixJ TCS. The aptamers motif 1 and 3 were interleaved with the ribosome-binding site of the *Ds*Red reporter gene in the pDusk and pDawn plasmids^25^, which additionally bore a PAL expression cassette. The resultant pDawn variants were further modified by introduction of the V28I exchange in the YF1 receptor which slows down dark recovery. Bacterial cultures harboring the plasmids were grown at 37°C in darkness (grey bars) or at two different blue-light intensities (5 and 60 μW cm^-2^, 470 nm, light blue and dark blue bars), and the *Ds*Red fluorescence was measured.

## References

(1) Möglich, A.; Yang, X.; Ayers, R. A.; Moffat, K. Structure and Function of Plant Photoreceptors. Annu. Rev. Plant Biol. 2010, 61, 21–47. https://doi.org/10.1146/annurev-arplant-042809-112259.

(2) Christie, J. M.; Reymond, P.; Powell, G. K.; Bernasconi, P.; Raibekas, A. A.; Liscum, E.; Briggs, W. R. Arabidopsis NPH1: A Flavoprotein with the Properties of a Photoreceptor for Phototropism. Science 1998, 282 (5394), 1698–1701.

(3) Salomon, M.; Eisenreich, W.; Dürr, H.; Schleicher, E.; Knieb, E.; Massey, V.; Rüdiger, W.; Müller, F.; Bacher, A.; Richter, G. An Optomechanical Transducer in the Blue Light Receptor Phototropin from Avena Sativa. Proc. Natl. Acad. Sci. 2001, 98 (22), 12357–12361. https://doi.org/10.1073/pnas.221455298.

(4) Crosson, S.; Moffat, K. Photoexcited Structure of a Plant Photoreceptor Domain Reveals a Light-Driven Molecular Switch. Plant Cell 2002, 14 (5), 1067–1075.

(5) Yee, E. F.; Diensthuber, R. P.; Vaidya, A. T.; Borbat, P. P.; Engelhard, C.; Freed, J. H.; Bittl, R.; Möglich, A.; Crane, B. R. Signal Transduction in Light-Oxygen-Voltage Receptors Lacking the Adduct-Forming Cysteine Residue. Nat. Commun. 2015, 6, 10079. https://doi.org/10.1038/ncomms10079.

(6) Dietler, J.; Gelfert, R.; Kaiser, J.; Borin, V.; Renzl, C.; Pilsl, S.; Ranzani, A. T.; García de Fuentes, A.; Gleichmann, T.; Diensthuber, R. P.; Weyand, M.; Mayer, G.; Schapiro, I.; Möglich, A. Signal Transduction in Light-Oxygen-Voltage Receptors Lacking the Active-Site Glutamine. Nat. Commun. 2022, 13 (1), 2618. https://doi.org/10.1038/s41467-022-30252-4.

(7) Crosson, S.; Rajagopal, S.; Moffat, K. The LOV Domain Family: Photoresponsive Signaling Modules Coupled to Diverse Output Domains. Biochemistry 2003, 42 (1), 2–10.

(8) Glantz, S. T.; Carpenter, E. J.; Melkonian, M.; Gardner, K. H.; Boyden, E. S.; Wong, G. K.-S.; Chow, B. Y. Functional and Topological Diversity of LOV Domain Photoreceptors. Proc. Natl. Acad. Sci. U. S. A. 2016, 113 (11), E1442–1451. https://doi.org/10.1073/pnas.1509428113.

(9) Alexandre, M. T. A.; Arents, J. C.; van Grondelle, R.; Hellingwerf, K. J.; Kennis, J. T. M. A Base-Catalyzed Mechanism for Dark State Recovery in the Avena Sativa Phototropin-1 LOV2 Domain. Biochemistry 2007, 46 (11), 3129–3137. https://doi.org/10.1021/bi062074e.

(10) Pudasaini, A.; El-Arab, K. K.; Zoltowski, B. D. LOV-Based Optogenetic Devices: Light-Driven Modules to Impart Photoregulated Control of Cellular Signaling. Front. Mol. Biosci. 2015, 2, 18. https://doi.org/10.3389/fmolb.2015.00018.

(11) Deisseroth, K.; Feng, G.; Majewska, A. K.; Miesenböck, G.; Ting, A.; Schnitzer, M. J. Next-Generation Optical Technologies for Illuminating Genetically Targeted Brain Circuits. J. Neurosci. 2006, 26 (41), 10380–10386. https://doi.org/10.1523/JNEUROSCI.3863-06.2006.

(12) Nagel, G.; Ollig, D.; Fuhrmann, M.; Kateriya, S.; Musti, A. M.; Bamberg, E.; Hegemann, P. Channelrhodopsin-1: A Light-Gated Proton Channel in Green Algae. Science 2002, 296 (5577), 2395–2398. https://doi.org/10.1126/science.1072068.

(13) Boyden, E. S.; Zhang, F.; Bamberg, E.; Nagel, G.; Deisseroth, K. Millisecond-Timescale, Genetically Targeted Optical Control of Neural Activity. Nat. Neurosci. 2005, 8 (9), 1263–1268. https://doi.org/10.1038/nn1525.

(14) Losi, A.; Gardner, K. H.; Möglich, A. Blue-Light Receptors for Optogenetics. Chem. Rev. 2018, 118 (21), 10659–10709. https://doi.org/10.1021/acs.chemrev.8b00163.

(15) Tang, K.; Beyer, H. M.; Zurbriggen, M. D.; Gärtner, W. The Red Edge: Bilin-Binding Photoreceptors as Optogenetic Tools and Fluorescence Reporters. Chem. Rev. 2021, 121 (24), 14906–14956. https://doi.org/10.1021/acs.chemrev.1c00194.

(16) Weber, A. M.; Kaiser, J.; Ziegler, T.; Pilsl, S.; Renzl, C.; Sixt, L.; Pietruschka, G.; Moniot, S.; Kakoti, A.; Juraschitz, M.; Schrottke, S.; Steegborn, C.; Bittl, R.; Mayer, G.; Möglich, A. A Blue Light Receptor That Mediates RNA Binding and Translational Regulation. Nat Chem Biol 2019, 15 (11), 1085–1092. https://doi.org/10.1038/s41589-019-0346-y.

(17) Renzl, C.; Kakoti, A.; Mayer, G. Aptamer-Mediated Reversible Transactivation of Gene Expression by Light. Angew. Chem. Int. Ed. 2020, 59 (50), 22414–22418. https://doi.org/10.1002/anie.202009240.

(18) Pilsl, S.; Morgan, C.; Choukeife, M.; Möglich, A.; Mayer, G. Optoribogenetic Control of Regulatory RNA Molecules. Nat. Commun. 2020, 11 (1), 4825. https://doi.org/10.1038/s41467-020-18673-5.

(19) Pietruschka, G.; Ranzani, A. T.; Weber, A. M.; Renzl, C.; Pilsl, S.; Patwari, T.; Möglich, A.; Mayer, G. Optozyme Mediated Control of Gene Expression. 2022.

(20) Liu, R.; Yang, J.; Yao, J.; Zhao, Z.; He, W.; Su, N.; Zhang, Z.; Zhang, C.; Zhang, Z.; Cai, H.; Zhu, L.; Zhao, Y.; Quan, S.; Chen, X.; Yang, Y. Optogenetic Control of RNA Function and Metabolism Using Engineered Light-Switchable RNA-Binding Proteins. Nat. Biotechnol. 2022, 1–8. https://doi.org/10.1038/s41587-021-01112-1.

(21) Murray, H. D.; Appleman, J. A.; Gourse, R. L. Regulation of the Escherichia Coli RrnB P2 Promoter. J. Bacteriol. 2003, 185 (1), 28–34. https://doi.org/10.1128/JB.185.1.28-34.2003.

(22) Deuschle, U.; Kammerer, W.; Gentz, R.; Bujard, H. Promoters of Escherichia Coli: A Hierarchy of in Vivo Strength Indicates Alternate Structures. EMBO J. 1986, 5 (11), 2987–2994.

(23) Salis, H. M.; Mirsky, E. A.; Voigt, C. A. Automated Design of Synthetic Ribosome Binding Sites to Control Protein Expression. Nat. Biotechnol. 2009, 27 (10), 946–950. https://doi.org/10.1038/nbt.1568.

(24) Lalwani, M. A.; Ip, S. S.; Carrasco-López, C.; Day, C.; Zhao, E. M.; Kawabe, H.; Avalos, J. L. Optogenetic Control of the Lac Operon for Bacterial Chemical and Protein Production. Nat. Chem. Biol. 2021, 17 (1), 71–79. https://doi.org/10.1038/s41589-020-0639-1.

(25) Ohlendorf, R.; Vidavski, R. R.; Eldar, A.; Moffat, K.; Möglich, A. From Dusk till Dawn: One-Plasmid Systems for Light-Regulated Gene Expression. J. Mol. Biol. 2012, 416 (4), 534–542. https://doi.org/10.1016/j.jmb.2012.01.001.

(26) Tabor, J. J.; Levskaya, A.; Voigt, C. A. Multichromatic Control of Gene Expression in Escherichia Coli. J. Mol. Biol. 2011, 405 (2), 315–324. https://doi.org/10.1016/j.jmb.2010.10.038.

(27) Multamäki, E.; García de Fuentes, A.; Sieryi, O.; Bykov, A.; Gerken, U.; Ranzani, A. T.; Köhler, J.; Meglinski, I.; Möglich, A.; Takala, H. Optogenetic Control of Bacterial Expression by Red Light. SSRN 2022. https://doi.org/10.2139/ssrn.4108992.

(28) Elowitz, M. B.; Leibler, S. A Synthetic Oscillatory Network of Transcriptional Regulators. Nature 2000, 403 (6767), 335–338. https://doi.org/10.1038/35002125.

(29) Stanton, B. C.; Nielsen, A. A. K.; Tamsir, A.; Clancy, K.; Peterson, T.; Voigt, C. A. Genomic Mining of Prokaryotic Repressors for Orthogonal Logic Gates. Nat. Chem. Biol. 2014, 10 (2), 99–105. https://doi.org/10.1038/nchembio.1411.

(30) Levskaya, A.; Chevalier, A. A.; Tabor, J. J.; Simpson, Z. B.; Lavery, L. A.; Levy, M.; Davidson, E. A.; Scouras, A.; Ellington, A. D.; Marcotte, E. M.; Voigt, C. A. Synthetic Biology: Engineering Escherichia Coli to See Light. Nature 2005, 438 (7067), 441–442. https://doi.org/10.1038/nature04405.

(31) Fernandez-Rodriguez, J.; Moser, F.; Song, M.; Voigt, C. A. Engineering RGB Color Vision into Escherichia Coli. Nat. Chem. Biol. 2017, 13 (7), 706–708. https://doi.org/10.1038/nchembio.2390.

(32) Jin, X.; Riedel-Kruse, I. H. Biofilm Lithography Enables High-Resolution Cell Patterning via Optogenetic Adhesin Expression. Proc. Natl. Acad. Sci. U. S. A. 2018, 115 (14), 3698–3703. https://doi.org/10.1073/pnas.1720676115.

(33) Farzadfard, F.; Lu, T. K. Synthetic Biology. Genomically Encoded Analog Memory with Precise in Vivo DNA Writing in Living Cell Populations. Science 2014, 346 (6211), 1256272. https://doi.org/10.1126/science.1256272.

(34) Tang, W.; Liu, D. R. Rewritable Multi-Event Analog Recording in Bacterial and Mammalian Cells. Science 2018, 360 (6385). https://doi.org/10.1126/science.aap8992.

(35) Cui, M.; Sun, T.; Li, S.; Pan, H.; Liu, J.; Zhang, X.; Li, L.; Li, S.; Wei, C.; Yu, C.; Yang, C.; Ma, N.; Ma, B.; Lu, S.; Chang, J.; Zhang, W.; Wang, H. NIR Light-Responsive Bacteria with Live Bio-Glue Coatings for Precise Colonization in the Gut. Cell Rep. 2021, 36 (11), 109690. https://doi.org/10.1016/j.celrep.2021.109690.

(36) Moser, F.; Tham, E.; González, L. M.; Lu, T. K.; Voigt, C. A. Light-Controlled, High-Resolution Patterning of Living Engineered Bacteria Onto Textiles, Ceramics, and Plastic. Adv. Funct. Mater. 2019, 29 (30), 1901788. https://doi.org/10.1002/adfm.201901788.

(37) Wang, Y.; An, B.; Xue, B.; Pu, J.; Zhang, X.; Huang, Y.; Yu, Y.; Cao, Y.; Zhong, C. Living Materials Fabricated via Gradient Mineralization of Light-Inducible Biofilms. Nat. Chem. Biol. 2021, 17 (3), 351–359. https://doi.org/10.1038/s41589-020-00697-z.

(38) Möglich, A.; Ayers, R. A.; Moffat, K. Design and Signaling Mechanism of Light-Regulated Histidine Kinases. J. Mol. Biol. 2009, 385 (5), 1433–1444. https://doi.org/10.1016/j.jmb.2008.12.017.

(39) Nihongaki, Y.; Suzuki, H.; Kawano, F.; Sato, M. Genetically Engineered Photoinducible Homodimerization System with Improved Dimer-Forming Efficiency. ACS Chem. Biol. 2014, 9 (3), 617–621. https://doi.org/10.1021/cb400836k.

(40) Hennemann, J.; Iwasaki, R. S.; Grund, T. N.; Diensthuber, R. P.; Richter, F.; Möglich, A. Optogenetic Control by Pulsed Illumination. Chembiochem 2018, 19 (12), 1296–1304. https://doi.org/10.1002/cbic.201800030.

(41) Ziegler, T.; Möglich, A. Photoreceptor Engineering. Front. Mol. Biosci. 2015, 2, 30. https://doi.org/10.3389/fmolb.2015.00030.

(42) Baumschlager, A.; Khammash, M. Synthetic Biological Approaches for Optogenetics and Tools for Transcriptional Light-Control in Bacteria. Adv. Biol. 2021, 5 (5), 2000256. https://doi.org/10.1002/adbi.202000256.

(43) Lindner, F.; Diepold, A. Optogenetics in Bacteria – Applications and Opportunities. FEMS Microbiol. Rev. 2022, 46 (2), fuab055. https://doi.org/10.1093/femsre/fuab055.

(44) Hoffman, S. M.; Tang, A. Y.; Avalos, J. L. Optogenetics Illuminates Applications in Microbial Engineering. Annu. Rev. Chem. Biomol. Eng. 2022, 13 (1), null. https://doi.org/10.1146/annurev-chembioeng-092120-092340.

(45) Hartsough, L. A.; Park, M.; Kotlajich, M. V.; Lazar, J. T.; Han, B.; Lin, C.-C. J.; Musteata, E.; Gambill, L.; Wang, M. C.; Tabor, J. J. Optogenetic Control of Gut Bacterial Metabolism to Promote Longevity. eLife 2020, 9, e56849. https://doi.org/10.7554/eLife.56849.

(46) Ong, N. T.; Tabor, J. J. A Miniaturized Escherichia Coli Green Light Sensor with High Dynamic Range. ChemBioChem 2018, 19 (12), 1255–1258. https://doi.org/10.1002/cbic.201800007.

(47) Chen, X.; Liu, R.; Ma, Z.; Xu, X.; Zhang, H.; Xu, J.; Ouyang, Q.; Yang, Y. An Extraordinary Stringent and Sensitive Light-Switchable Gene Expression System for Bacterial Cells. Cell Res. 2016, 26 (7), 854–857. https://doi.org/10.1038/cr.2016.74.

(48) Li, X.; Zhang, C.; Xu, X.; Miao, J.; Yao, J.; Liu, R.; Zhao, Y.; Chen, X.; Yang, Y. A Single-Component Light Sensor System Allows Highly Tunable and Direct Activation of Gene Expression in Bacterial Cells. Nucleic Acids Res. 2020, 48 (6), e33–e33. https://doi.org/10.1093/nar/gkaa044.

(49) Dietler, J.; Schubert, R.; Krafft, T. G. A.; Meiler, S.; Kainrath, S.; Richter, F.; Schweimer, K.; Weyand, M.; Janovjak, H.; Möglich, A. A Light-Oxygen-Voltage Receptor Integrates Light and Temperature. J. Mol. Biol. 2021, 433 (15), 167107. https://doi.org/10.1016/j.jmb.2021.167107.

(50) Ryu, M.-H.; Gomelsky, M. Near-Infrared Light Responsive Synthetic c-Di-GMP Module for Optogenetic Applications. ACS Synth. Biol. 2014, 3 (11), 802–810. https://doi.org/10.1021/sb400182x.

(51) Lynch, S. A.; Desai, S. K.; Sajja, H. K.; Gallivan, J. P. A High-Throughput Screen for Synthetic Riboswitches Reveals Mechanistic Insights into Their Function. Chem. Biol. 2007, 14 (2), 173–184. https://doi.org/10.1016/j.chembiol.2006.12.008.

(52) de Jesus, V.; Qureshi, N. S.; Warhaut, S.; Bains, J. K.; Dietz, M. S.; Heilemann, M.; Schwalbe, H.; Fürtig, B. Switching at the Ribosome: Riboswitches Need RProteins as Modulators to Regulate Translation. Nat. Commun. 2021, 12 (1), 4723. https://doi.org/10.1038/s41467-021-25024-5.

(53) Weigand, J. E.; Sanchez, M.; Gunnesch, E.-B.; Zeiher, S.; Schroeder, R.; Suess, B. Screening for Engineered Neomycin Riboswitches That Control Translation Initiation. RNA 2008, 14 (1), 89–97. https://doi.org/10.1261/rna.772408.

(54) Breaker, R. R. Riboswitches and Translation Control. Cold Spring Harb. Perspect. Biol. 2018, 10 (11), a032797. https://doi.org/10.1101/cshperspect.a032797.

(55) Strack, R. L.; Strongin, D. E.; Bhattacharyya, D.; Tao, W.; Berman, A.; Broxmeyer, H. E.; Keenan, R. J.; Glick, B. S. A Noncytotoxic DsRed Variant for Whole-Cell Labeling. Nat. Methods 2008, 5 (11), 955–957. https://doi.org/10.1038/nmeth.1264.

(56) Gibson, D. G.; Young, L.; Chuang, R.-Y.; Venter, J. C.; Hutchison, C. A.; Smith, H. O. Enzymatic Assembly of DNA Molecules up to Several Hundred Kilobases. Nat. Methods 2009, 6 (5), 343–345. https://doi.org/10.1038/nmeth.1318.

(57) Mathes, T.; Vogl, C.; Stolz, J.; Hegemann, P. In Vivo Generation of Flavoproteins with Modified Cofactors. J. Mol. Biol. 2009, 385 (5), 1511–1518. https://doi.org/10.1016/j.jmb.2008.11.001.

(58) Möglich, A. An Open-Source, Cross-Platform Resource for Nonlinear Least-Squares Curve Fitting. J. Chem. Educ. 2018, 95 (12), 2273–2278. https://doi.org/10.1021/acs.jchemed.8b00649.

